# Neuronal control of maternal provisioning in response to social cues

**DOI:** 10.1101/2021.02.01.429208

**Authors:** Jadiel A. Wasson, Gareth Harris, Sabine Keppler-Ross, Trisha J. Brock, Abdul R. Dar, Rebecca A. Butcher, Sylvia E.J. Fischer, Konstantinos Kagias, Jon Clardy, Yun Zhang, Susan Mango

## Abstract

Mothers contribute cytoplasmic components to their progeny in a process called maternal provisioning. Provisioning is influenced by the parental environment, but the molecular pathways that transmit environmental cues from mother to progeny are not well understood. Here we show that in *C. elegans*, social cues modulate maternal provisioning to regulate gene silencing in offspring. Intergenerational signal transmission depends on a pheromone-sensing neuron and neuronal FMRF (Phe-Met-Arg-Phe)-like peptides. Parental FMRF signaling promotes the deposition of mRNAs for translational components in progeny, which in turn reduces gene silencing. Previous studies had implicated FMRF signaling in short-term responses such as modulated feeding behavior in response to the metabolic state^1,2^, but our data reveal a broader role, to coordinate energetically expensive processes such as translation and maternal provisioning. This study identifies a new pathway for intergenerational communication, distinct from previously discovered pathways involving small RNAs and chromatin, that links sensory perception to maternal provisioning.

---

We sought to identify factors involved in communication from the parental environment to progeny in *C. elegans*. Environmentally supplied double-stranded RNA (dsRNA) induces gene silencing in the next generation, providing a useful assay^3^. We fed mothers diluted dsRNA for the essential gene *pha-4*, which generated weak silencing in wild-type animals, and screened for mutants that enhanced RNAi potency. Silencing was monitored as lethality after *pha-4* RNAi (Figure 1a) or reduced PHA-4 expression (see below). We found that the strength of RNAi was enhanced in mutants that were defective in pheromone biosynthesis: mutations in *daf-22/SCP2, dhs-28/HSD*, or *acox-1.1/ACOX1*^4–6^ each generated more lethality compared to wild-type worms (Figure 1b). Addition of conditioned media made from wild-type animals restored the RNAi response of *daf-22* mutants, whereas conditioned media made from *daf-22* mutants did not (Figure 1b). This result indicates that RNAi in progeny responds to molecules synthesized by the *daf-22* pathway and secreted into the environment. Moreover, exposure to conditioned media for 24 hours during the last larval stage and early adulthood was sufficient for rescue, indicating that early life exposure of parents to the environment was not necessary. The *daf-22* pathway generates multiple pheromones that signal the presence of conspecifics, including ascarosides asc-wC3, asc-C6-MK, and asc-DC9, which induce a developmental arrest (dauer)^5–7^. However, a synthetic mixture of these three dauer pheromones, which induced 95% dauers under established conditions (data not shown), did not rescue the enhanced RNAi of *daf-22* mutants^5,6^. These data suggest that RNAi in progeny is modulated by exposure of mothers to conspecific pheromones in the environment, and these are distinct from dauer pheromones.

**Figure 1:**
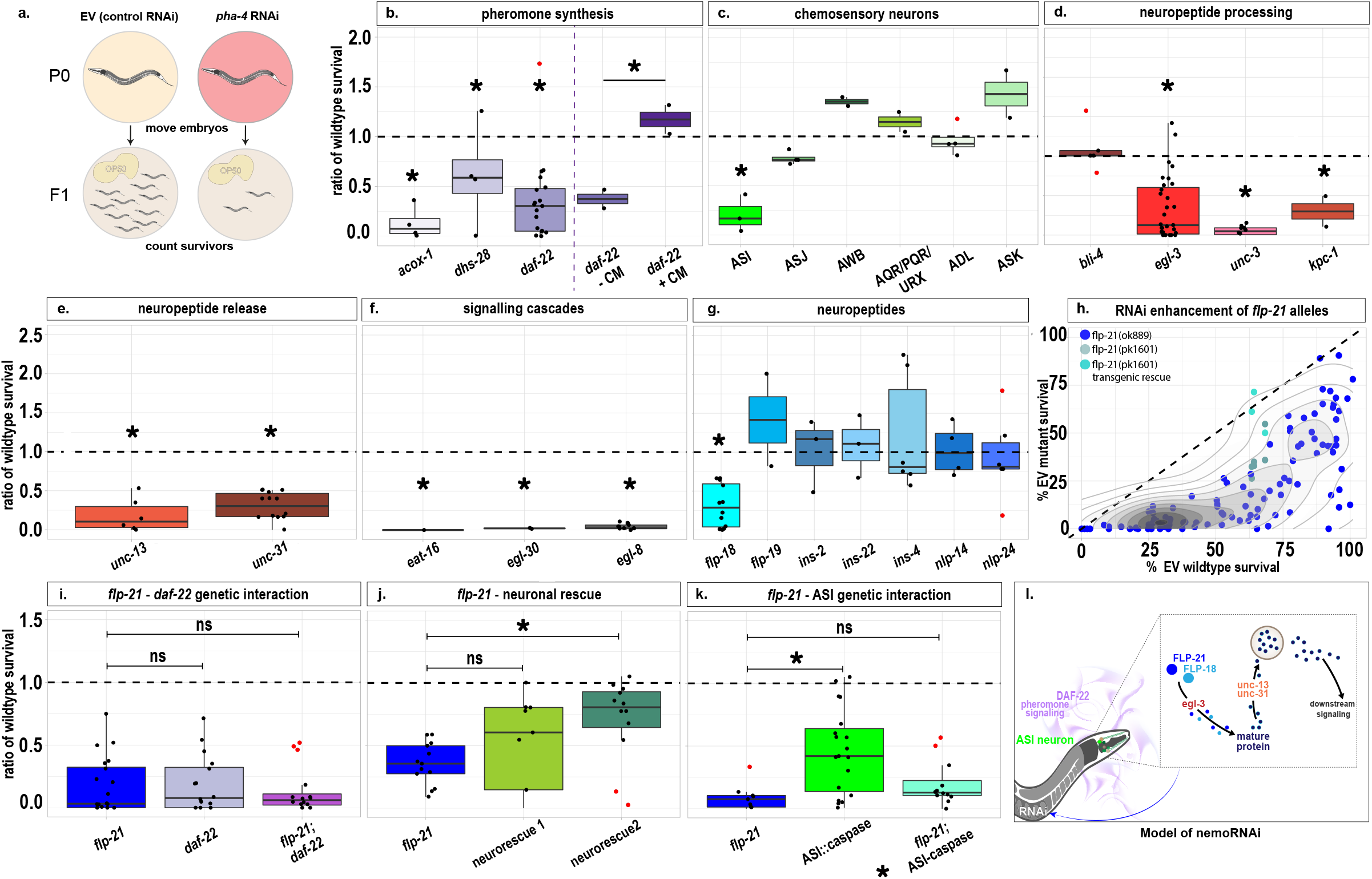
Neuronal signaling modulates RNAi. (a) Schematic of *pha-4* RNAi survival experiments. Survey for enhanced RNAi by genes involved in (b) pheromone synthesis, (c) chemosensory neurons, (d) neuropeptide processing, (e) neuropeptide release, (f) signalling factors, (g) neuropeptides, (h) *flp-21*. Enhanced RNAi is rescued by (b) conditioned media (CM) from wild-type animals but not from *daf-22* mutants, (h) a genomic transgene for *flp-21* for *flp-21(pk1601)* or (j) a neuronally expressed *flp-21* for *flp-21(ok889)*. Epistasis analysis between *flp-21* and (i) *daf-22* or (k) loss of ASI generated by caspase. (l) A model for nemoRNAi. Data visualization represents the ratio of wild-type %EV survival for each mutant (wt represented by a dotted line at 1 on each graph). Boxplots indicate the first quartile (Q1) to third quartile (Q3) with box, median and quartile 2 of data indicated by a horizontal line, whiskers go from Q1 to smallest non-outlier and from Q3 to largest non-outlier. Additional outliers plotted as red dots outside whiskers. Each boxplot represents at least 3 biological replicates. Individual experiments represented as points on boxplots. *p < 0.05. The details of the data analysis are included in Methods.

Small molecule pheromones are detected by chemosensory neurons, including ADL, ASK and ASI^8–14^. We discovered that killing the ASI neurons enhanced RNAi in the progeny, whereas inactivation of ADL, ASK or several other sensory neurons did not (Figure 1c). In addition, inactivating *unc-3*, a transcription factor required for ASI cell fate^15^, also enhanced *pha-4* RNAi (Figure 1d). Consistent with the idea that ASI-mediated signaling regulates RNAi in progeny, genes encoding neuronal signaling components modulated RNAi. *egl-3* and *kpc-1*, which encode pro-protein convertases that process signaling peptides^16,17^, both enhanced RNAi when inactivated, whereas the endo-protease *bli-4* did not^18^ (Figure 1d). Signaling peptides depend on calciumdependent activator proteins (CAPS)^19,20^ for release from dense core vesicles, and the CAPS factor *unc-31* enhanced RNAi when mutated. Similarly, *unc-13*, which is required for peptide and vesicle release during neurotransmission^21^, also enhanced RNAi when mutated (Figure 1e).

The requirement for the ASI neuron, as well as *egl-3, kpc-1, unc-13* and *unc-31*, implicated neuropeptide-encoding genes for a role in maternal modulation of RNAi. Neuropeptides fall into three categories, the FMRF-amide peptides (FLPs), the insulinlike peptides (ILPs), and the neuropeptide-like peptides (NLPs)^22^. We surveyed candidates from all three categories and discovered that two FLP-like genes, *flp-18* and *flp-21*^1,2^, were involved in modulating RNAi in progeny. Two mutant alleles each of *flp-18* and *flp-21* enhanced RNAi (Figure 1g, h), and did so over a range of dsRNA dilutions (Figures 1h and 2a). Expressing a genomic fragment containing the wild-type *flp-21* gene rescued the enhanced RNAi phenotype (Figure 1h). In contrast, inactivation of several ILPs or NLPs showed no effect, even though insulin signaling has been implicated in other examples of intergenerational communication^23–25^.

**Figure 2.**
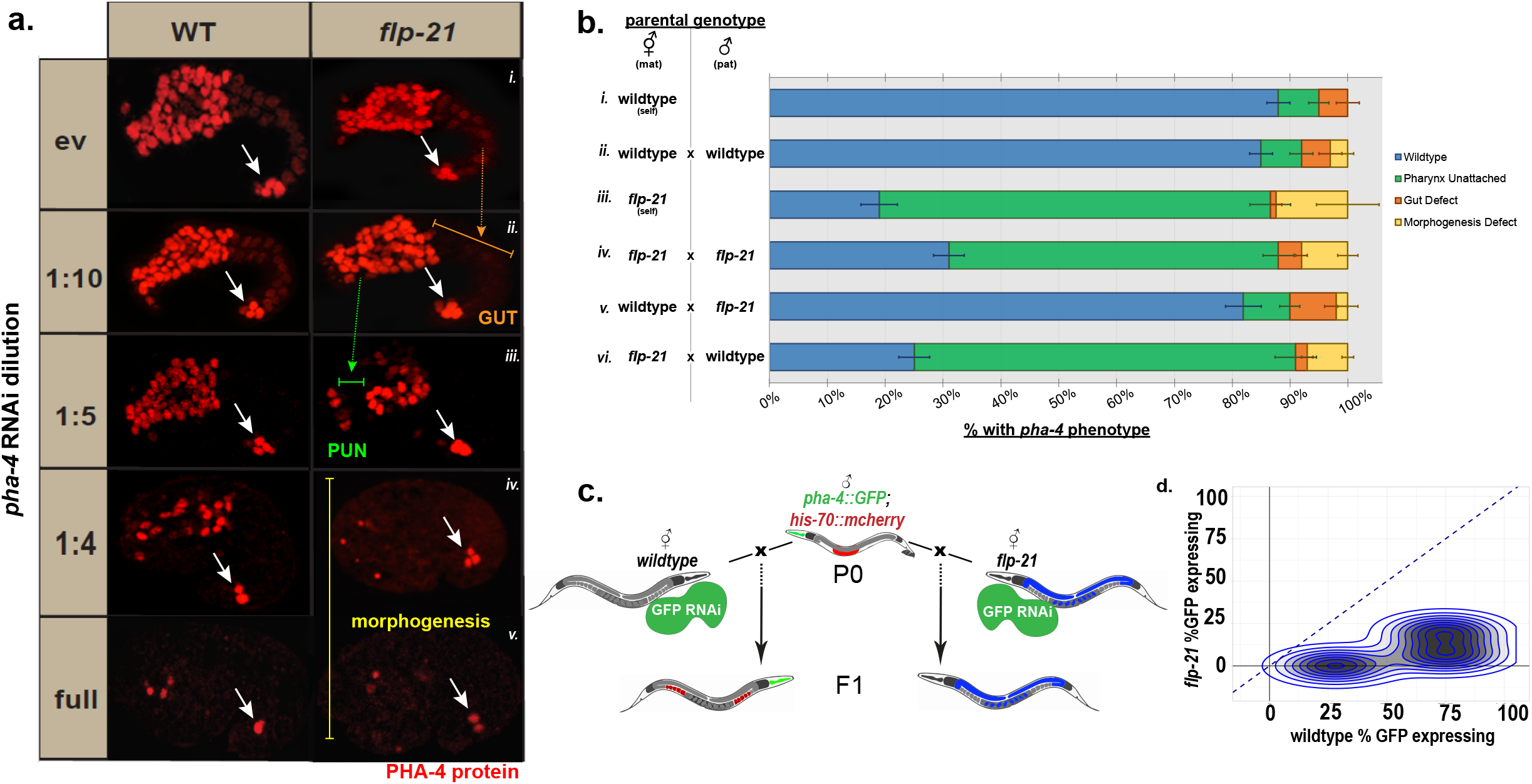
nemoRNAi is maternal and does not require the target locus. (a) PHA-4 protein in wild-type (WT) or *flp-21* embryos treated with varying dilutions of dsRNA. Phenotypes of gut defects, pharynx unattached (Pun), and morphogenesis denoted by orange, green and yellow, respectively. Note the hindgut stains in all samples, a positive staining control since it is refractory to RNAi (arrow). (b) Self progeny (i and iii) or progeny from mating (ii, iv, v, vi) were examined for *pha-4* phenotypes after RNAi. Mother (mat) and father (pat). Quantitation of *pha-4* phenotypes (wildtype in blue, Pun in green, gut defective in orange, and morphogenesis defect in yellow) in progeny from each of the indicated crosses reveals that flp-21 mutation is required in mothers. Most dramatic phenotype is graphed. Error bars indicate standard deviation across three biological replicates. (c) Schematic of cross to determine if target DNA is required in mothers for enhancement. Wildtype or *flp-21* mutants lacking a *gfp* transgene are exposed to *gfp* dsRNA, then mated to *pha-4::gfp* males (with linked *his-7O::mcherry* to identify cross progeny). (d) Graph of data from (c) indicating percent progeny expressing GFP from wild-type (x-axis) or *flp-21* (y-axis) mothers (each point is the average GFP expression from total number of progeny in one paired experiment, 3 biological replicates performed. Dotted line represents 1:1 GFP expression in wildtype vs *flp-21* progeny. Wilcoxon rank sum, *p<0.05.

RNAi enhancement was rescued by a single-copy transgene expressing *flp-21* in neurons (Figure 1j). We used a promoter expressed broadly in the nervous system and tested two independently generated lines. One rescued the enhanced RNAi phenotype to an almost wild-type level, and one rescued to an intermediate level, likely due to variable expression of the promoter in neurons^26^. These data suggest that *flp-21-* mediated signaling in neurons can modulate the strength of RNAi in the next generation.

To investigate the extent and gene-specificity of *flp-21* beyond *pha-4* RNAi, we surveyed dsRNAs targeting a variety of genes. *flp-21* mutants exhibited an enhanced RNAi response to a range of dsRNAs, including those targeting maternally-expressed genes (*mex-3, skn-1*), and genes activated during the first half of embryonic development (*dlg-1, par-3, hmr-1, pha-4*) (Figure 2a, Supplemental Figure 1a). However, two genes expressed during terminal embryogenesis were resistant to RNAi and failed to be enhanced by *flp-21* (*ifb-2* and *myo-2*) (Supplemental Figure 1a). Likewise, terminal *pha-4* expressed in the hindgut was refractory to RNAi, presumably because it is activated late (e.g. Figure 2a). We also found that *flp-21* mutants enhanced RNAi over a range of dsRNA exposure, from 18 to 43 hours (Supplemental Figure 1b). These data indicate that loss of *flp-21* results in a generally enhanced RNA interference (‘Eri’) phenotype for genes expressed during early embryo development. We call this Eri phenotype neuronal enhancement by mother of RNAi (‘nemoRNAi’, Figure 1l).

FMRF-amide related neuropeptides signal to target tissues via G-protein coupled receptor signaling cascades^27,28^. We found that EGL-30, the Gq-protein subunit α^29^, induced a nemoRNAi phenotype when inactivated (Figure 1f). Furthermore, mutations in *eat-16*, the regulator of G-protein signaling, and *egl-8/PLC□4*, additional signaling components of Gαq pathways^29,30^, both had nemoRNAi phenotypes when exposed to diluted *pha-4* dsRNA (Figure 1f). Taken together, these results show that FMRF-amide related neuropeptides FLP-18, FLP-21 and Gαq signaling play a role in nemoRNAi.

Epistasis analysis between *flp-21* and *daf-22* indicated that these two factors act in the same pathway because double mutants did not produce an additive effect (Figure 1i). Similarly, *flp-21* mutations combined with ASI inactivation indicated that these also act in the same pathway (Figure 1k). These data suggest that a pheromone-responsive, FLP-specific chemosensory signaling pathway modulates gene silencing in progeny via RNAi (Figure 1l). We conclude that sensory perception, including that of social information, modulates silencing in progeny. We focus on *flp-21* for the remainder of the study, given its robust phenotype and lack of pleiotropy.

If FLP-21 links pheromone sensing to gene silencing in progeny, it predicts that FLP-21 functions in parents, not progeny. To test this idea, we examined the phenotypes of heterozygous F1 progeny generated by mating mutant and wild-type parents. Mutant *flp-21* mothers mated with wild-type fathers produced progeny that behaved like *flp-21* homozygotes (compare Figure 2a row vi with rows iii and iv), whereas wild-type mothers mated with mutant fathers produced offspring that behaved like wild-type (compare Figure 2a row v with rows I and ii). These data reveal that *flp-21* functions in mothers to modulate the RNAi response in progeny. We note that *flp-21* is expressed in the soma, and is not contributed to the embryo from the mothers’ germ lines^31^. These data support the hypothesis that the FLP-21-mediated signaling pathway acts in mothers to modulate RNAi in progeny (see discussion).

Given that chromatin modifications have been implicated in communication between mothers and offspring^32^, we tested whether the target gene of nemoRNAi had to be present in mothers, a condition required for chromatin modification. Wild-type or *flp-21* mutant mothers were exposed to dsRNA against *gfp* and mated to males bearing a *gfp* transgene (Figure 2c). The resulting progeny carried the paternal *gfp* transgene and a linked *histone::mCherry* control (Figure 2c). *flp-21* mutations enhanced *gfp* silencing in progeny, revealing that the target *gfp* DNA was not required in mothers to initiate nemoRNAi (Figure 2d). We conclude that chromatin modifications at the target locus are not involved in *flp-21-dependent* intergenerational communication. We recognize that indirect effects of chromatin modifications may still be involved.

As nemoRNAi did not require the target locus in mothers, a diffusible molecule such as RNA was an attractive candidate to be the vehicle of inheritance. We focused on mRNAs, as maternal deposition of mRNA regulates many aspects of embryogenesis^33,34^, and because we did not identify any small RNAs that could account for nemoRNAi (Supplemental Figure 2). We analyzed our mRNA sequencing libraries by differential expression analysis, which identified 3791 significantly deregulated genes in *flp-21* early embryos compared to wild-type (Figure 3a). These genes were enriched for translation/ribosomal components according to four categories of GO term analysis: Biological Pathway, Cell Component, Molecular Function, and KEGG Pathway (Figure 3b). Moreover, two ribosomal RNAs, *rrn-3.1* and *rrn-1.2*, and the elongation factor *eef-1A. 1* were the top drivers distinguishing wild-type from *flp-21* RNA samples by two-dimensional principal component analysis (PCA; Figure 3c). Additional differences between our RNA samples included vitellogenins (PCA), innate immunity (GO analysis) and protein phosphorylation (GO analysis), but only translational components were identified as top hits by both GO analysis and PCA.

**Figure 3.**
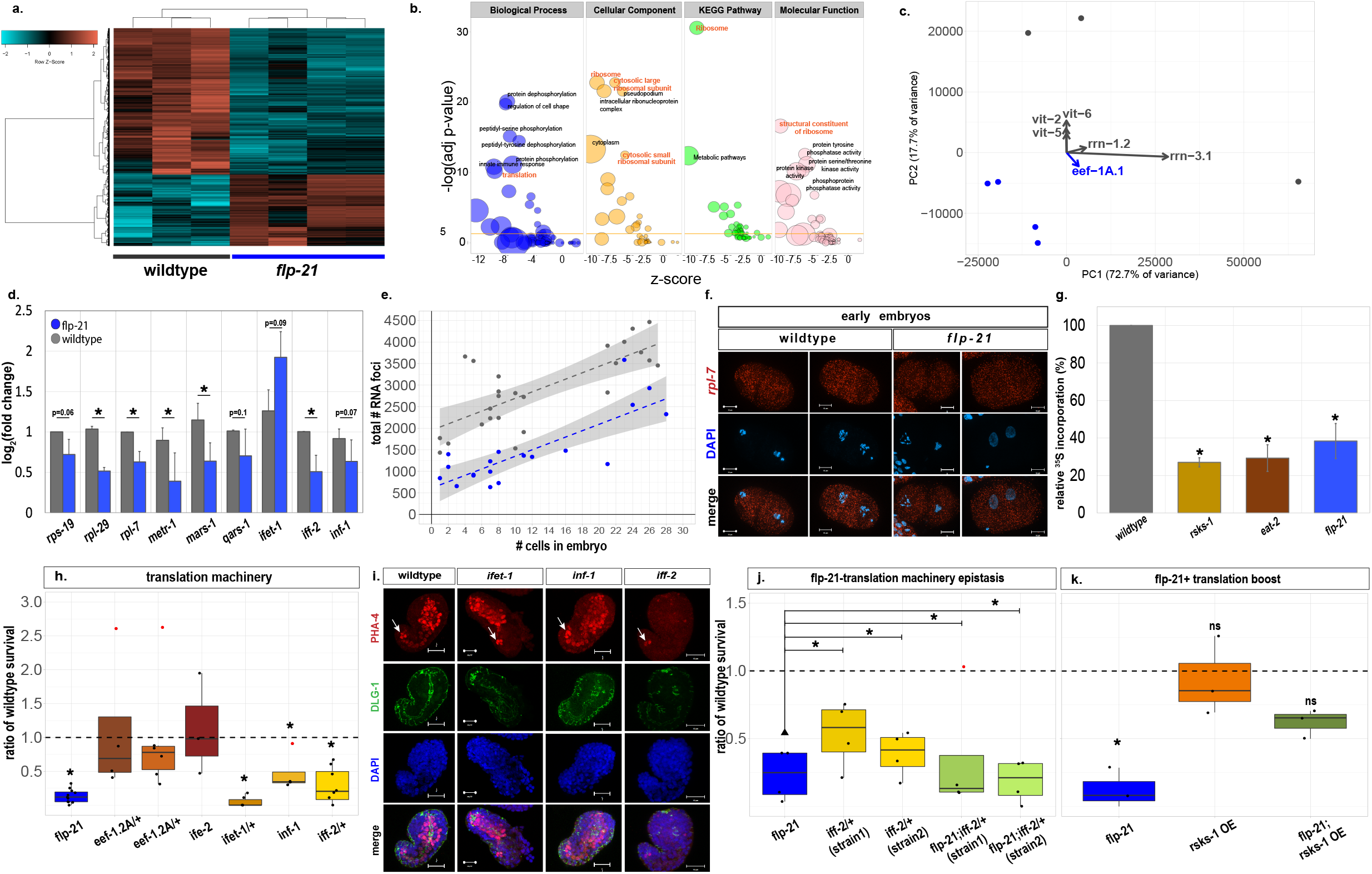
Translation is reduced in *flp-21* mutants and sufficient for nemoRNAi. (a) Heat map and hierarchical clustering of significant transcriptional changes revealed by DESEQ2 for three wild-type and four *flp-21* replicates (red upregulated, turquoise downregulated genes, z-scores indicated). (b) Bubble plots for GO analysis enriched terms identified with DAVID (significant genes from a). Four different categories of GO analysis and KEGG pathways indicated. Size of bubbles indicate the number of genes. X-axis indicates z score, y-axis indicates negative logarithm of adjusted p-value from GO analysis (higher = more significant). GO analysis enriched terms with log(adjusted p-value) < 10 labelled with ID. Translation and Ribosome are top hits in each GO category and in KEGG pathways. (c) PCA analysis of wild-type (grey circles) and flp-21 (blue circles) samples. Top genes driving clustering of samples labeled, and arrows indicate direction of driving change. PC1 on x-axis, PC2 on y-axis. (d) qPCR confirmation of select translational machinery components. 2 samples from each genotype were assayed in 3 independent experiments. Data presented as log2(foldchange). Statistics performed on ddCt values, ANOVA Tukey’s, *p<0.05. (e) Quantitation of smFISH for wild-type and *flp-21* mutants. X-axis represents embryonic age (# of cells), y-axis represents number of smFISH foci. (f) Representative smFISH (red) images for *rpl-7* (scale bars = 10uM). (g) Relative 35S incorporation. Wildtype set to 100%. *flp-21* mutants have similar incorporation rates as positive controls *rsks-1* and *eat-2*. ANOVA Tukey’s, *p<0.05. (h) *pha-4* RNAi survival assay performed on translation machinery genes. Data visualization represents ratio of wildtype (WT) %Empty Vector (EV) survival for each mutant (WT represented with dotted line at 1 on each graph). Three mutants (*inf-1, ifet-1*, and *iff-2*) in addition to *flp-21*, have an enhanced RNAi phenotype. Note hindgut staining (arrow). (i) Representative images of PHA-4 staining after *pha-4* RNAi for each genotype (scale bars = 10uM). Note the hindgut stains in all samples, a positive staining control since it is refractory to RNAi. (j) Genetic interaction between *flp-21* and *iff-2* heterozygote reveals that they are in the same pathway. (k) Genetic interaction between *flp-21* and *rsks-1* overexpression transgene rescues enhanced RNAi phenotype. ANOVA Dunnett’s test, *p<0.05.

We generated circle plots for the top 8 GO pathways to identify the directionality of the RNA changes within each GO category (Supplemental Figure 3). In each category, there was a robust downregulation of translation-associated terms. These results were confirmed by qPCR analysis for eight translation machinery genes spanning initiation, elongation etc. (Figure 3d); a ninth gene, *ifet-1*, was upregulated in *flp-21* mutants and encodes a translation inhibitor^35^ (Figure 3d). In addition, smFISH for ribosomal protein *rpl-7* revealed an almost two-fold decrease in *flp-21* mutants (Figure 3e, f). Taken together, these data indicate that many maternally provided translation components are reduced in *flp-21* early embryos.

The reduction of translation components in *flp-21* early embryos suggested a decrease in translation rate. To test this idea, we analyzed translation in wild-type and *flp-21* mutants with ^35^S metabolic labeling. ^35^S incorporation was reduced to less than 50% in *flp-21* mutants compared to wild-type animals, which was comparable to well-known translation-deficient mutants *rsks-1* and *eat-2*^36^ (Figure 3g). This result revealed that the ongoing translation rate in *flp-21* mutants was significantly reduced.

We next asked whether the translational changes of *flp-21* mutants could account for nemoRNAi. As many genes associated with translation are embryonic lethal, we relied on heterozygous mutations for some genes and surveyed these by *pha-4* RNAi. From our panel of mutants, three had enhanced RNAi phenotypes: *inf-1, ifet-1*, and *iff-2* (Figure 3h). Of these, *iff-2/+* mothers generated a reduction of PHA-4 staining in progeny that closely resembled *flp-21* mutants (Figure 3i). *inf-1* and *ifet-1* were both sickly, even for the EV control, so a possible role was less clear for these genes (Supplemental Figure 3b). These data suggest that not all translational components can modulate RNAi efficiency, but that reduction of *iff-2* is particularly effective for enhancing RNAi. *iff-2* encodes the orthologue of mammalian EIF5A and EIF5A2^37^. EIF5A is important both for translational elongation, particularly at ribosome stalling sites, and for termination^38^. EIF5A2 may also function in mRNA decay in response to stress^39^, and both factors have been implicated in human diseases such as cancer^40^.

If the reduction in translation is sufficient to explain nemoRNAi in *flp-21* mutants, then a combination of mutations should not have an additive effect on lethality associated with *pha-4* RNAi. This prediction was borne out*: flp-21; iff-2/+* double mutants resembled *flp-21* single mutants for the response to RNAi (Figure 3j). We analyzed two independently constructed strains of *flp-21; iff-2/+* along with single mutant strains for *iff-2/+* (two strains) and *flp-21* (one strain) generated from the siblings during the strain construction. The combination of *flp-21* mutations with a heterozygous *iff-2* allele resulted in no additive effect on RNAi lethality, indicating that these two genes are in the same pathway for nemoRNAi.

Pathways that regulate translation converge on TOR components and S6 kinase signaling, which includes LET-363 and RSKS-1 in *C. elegans*^41^. To determine if translation is key for nemoRNAi by *flp-21* mutations, we introduced a *rsks-1* overexpression transgene into the *flp-21* mutant background. Augmenting translation via *rsks-1* overexpression^42,43^ was able to rescue the enhanced RNAi phenotype of *flp-21* progeny (Figure 3k). As we have demonstrated that these maternal chemosensory pathways interact with, and respond to, environmental cues, this work suggests a model where the maternal environment can modulate embryonic gene silencing and development via provisioning of translation components.

In summary, our results indicate that *flp-21*-dependent signaling of social cues increases the maternal provisioning of translational components and reduces RNAi responsiveness in the offspring. Key supporting evidence for this model includes that the absence of social cues, chemosensory neurons or FMRF *flp-21* signaling increases RNAi in offspring. Loss of maternal *flp-21* signaling also leads to reduced translation and lower translation components in offspring. Our analyses indicate that *flp-21* is upstream of maternal provisioning of translational components, which in turn is upstream of RNAi responsiveness. While published studies had previously implicated FMRF signaling in short-term behaviors such as feeding^1,2^, our data reveal a broader role, to coordinate energetically expensive processes such as translation and maternal provisioning.

Many animals exhibit intergenerational communication regarding the parental environment, including humans, mice, *Drosophila* as well as *C. elegans*^44,45^. This study describes how manipulating an animal’s perception of its environment is sufficient to alter maternal provisioning of translation factors. This result contrasts with other studies where altered maternal provisioning occurs in response to physiological changes. For example, maternal provisioning of RNAs for translation components is reduced when *Drosophila* mothers are deprived of food^46^. However, in this example, it is unclear whether the reduction of translation components in progeny reflects dysregulation in starved mothers^47^ or an altered sensory response. Our results argue that sensory stimulation alone is sufficient, since *flp-21* mutants feed and grow as well as wild-type animals (Supplemental Figure 1c), yet they reduce their translational capacity. Reduction in *flp-21* signaling phenocopies and works in the same pathway as pheromone deprivation. Lack of pheromone signaling reflects the absence of conspecifics and may thereby indicate an adverse environment^4–7^. It is conceivable that mothers that sense this suboptimal environment protect their offspring by reducing translation (to conserve energy) and augmenting RNAi (to boost immunity). Similar processes may occur in other animals. For example, in rodents, mothers’ social experience is transmitted to progeny, but the mechanism is unknown^48^. Our results suggest that material provisioning underlies the transmission of social information across generations.

## Acknowledgements

We thank A. Schier and the Mango lab for comments on the manuscript, R. Centko for conditioned media, G. Fucile and X. Canales for informatics advice, the Biozentrum Imaging Core Facility (IMCF) and Quantitative Sequencing Facility for technical assistance. Strains were provided by the CGC (NIH grant P40 OD010440). This work was supported by the Helen Hay Whitney foundation to JAW; SNF 310030_185157 and NIH 5R37 GM056264 to SEM; Harvard University to YZ; NIH R01AT009874 to JC; NIH GM118775 and NSF Career-1555050 to RAB.

---

**Figure Supplement 1:**
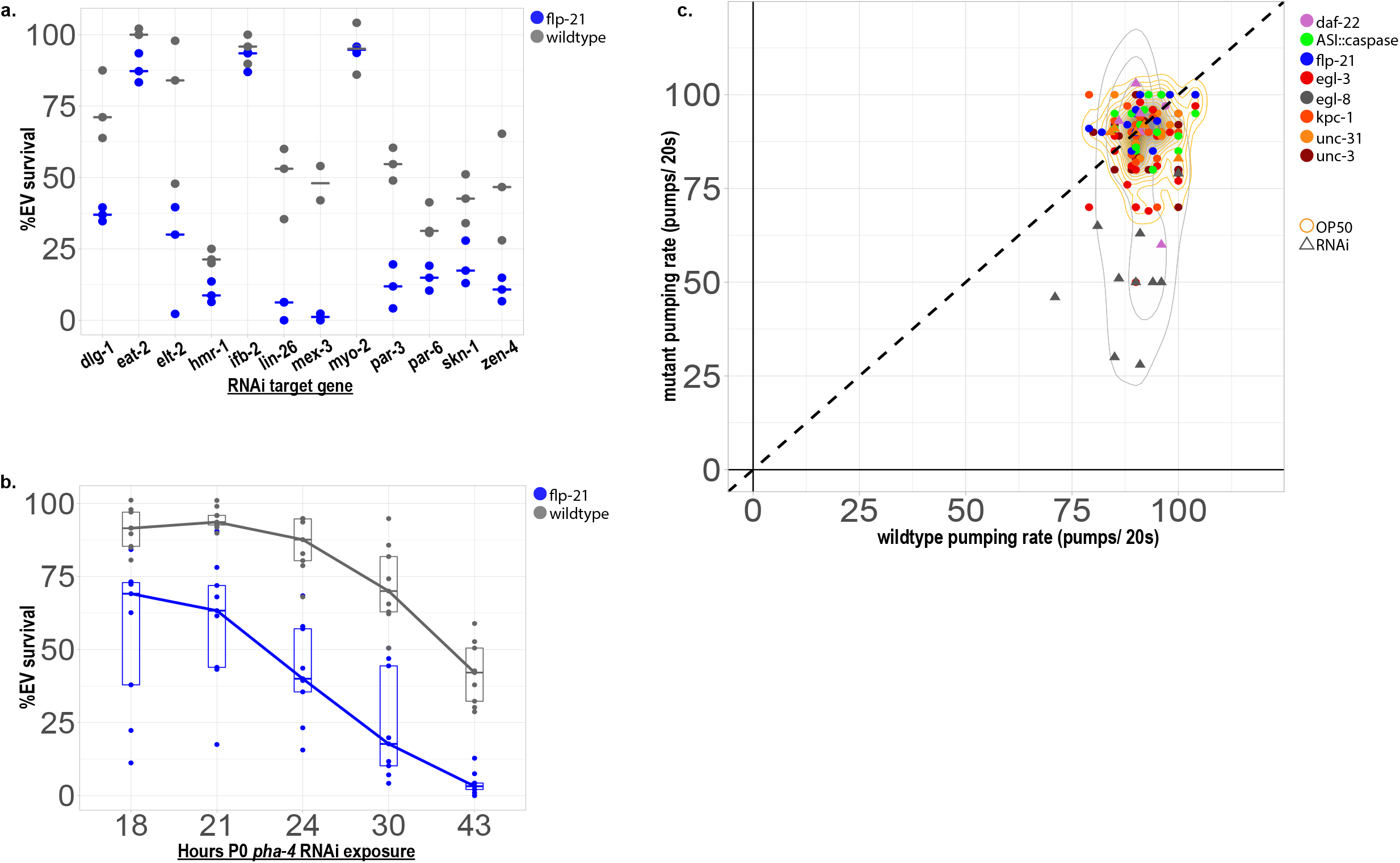
Characterization of the nemoRNAi effect. (a) RNAi screen on *flp-21* mutants targeting various developmentally important genes (x-axis). Y-axis indicates %EV survival. Three independent experiments performed with 100 embryos per genotype per experiment. Wilcoxon Rank Sum, *p<0.05. (b) Tracking of RNAi enhancement effect on progeny survival over 18, 21, 24, 30, and 43 hours of pha-4 RNAi exposure in P0 mothers. Three independent experiments performed with 100 embryos per genotype per timepoint per experiment. Wilcoxon Rank Sum, *p<0.05. (c) Pumping rates in various nemoRNAi mutants compared to wild-type pumping rates on OP50 (circles, orange density) or after RNAi exposure (triangles, grey density).

In mammalian systems, RNAs have been implicated as modulators of intergenerational phenotypes such as altered metabolism in high fat diets and neuronal changes in response to fear^1^–^3^. Small RNAs such as piRNAs and siRNAs in C. elegans, and miRNAs and tRFs in mammals regulate multi-generational effects^4^–^7^. We sequenced small RNAs (16 and 30 nt) and mRNAs from early embryos (1-10 cells, Supplemental Figure 2a). MicroRNA and rRNA class distributions compared to other small RNA classes were altered in our analysis (Supplemental Figure 2b-d). We further analyzed these changes with Taqman assays, but did not observe dramatic changes (Supplemental Figure 2e). We also interrogated the *pha-4* RNAi phenotypes for three top deregulated microRNAs, *miR-35, miR-51*, and *mir-72* (Supplemental Figure 2f). *miR-35* was the only microRNA tested that had a strong *pha-4* RNAi phenotype, which had been observed previously^8^. Epistasis analysis between flp-21 and *miR-35* loss-of-function revealed that their genetic interaction was additive, indicating that they work in parallel to affect RNAi (Supplemental Figure 2g). Moreover, *miR-35* was over expressed in *flp-21* mutants. In short, we were unable to identify microRNA changes that could account for nemoRNAi. Futhermore, small RNAs that mapped to genes (22G and 26G siRNAs), piRNAs, endogenous siRNAs, and transposons were largely unchanged between wild-type and *flp-21* embryos (Supplemental Figure 2b, h). These results suggested that alternative effector molecules were r esponsible for nemoRNAi.

**Figure Supplement 2:**
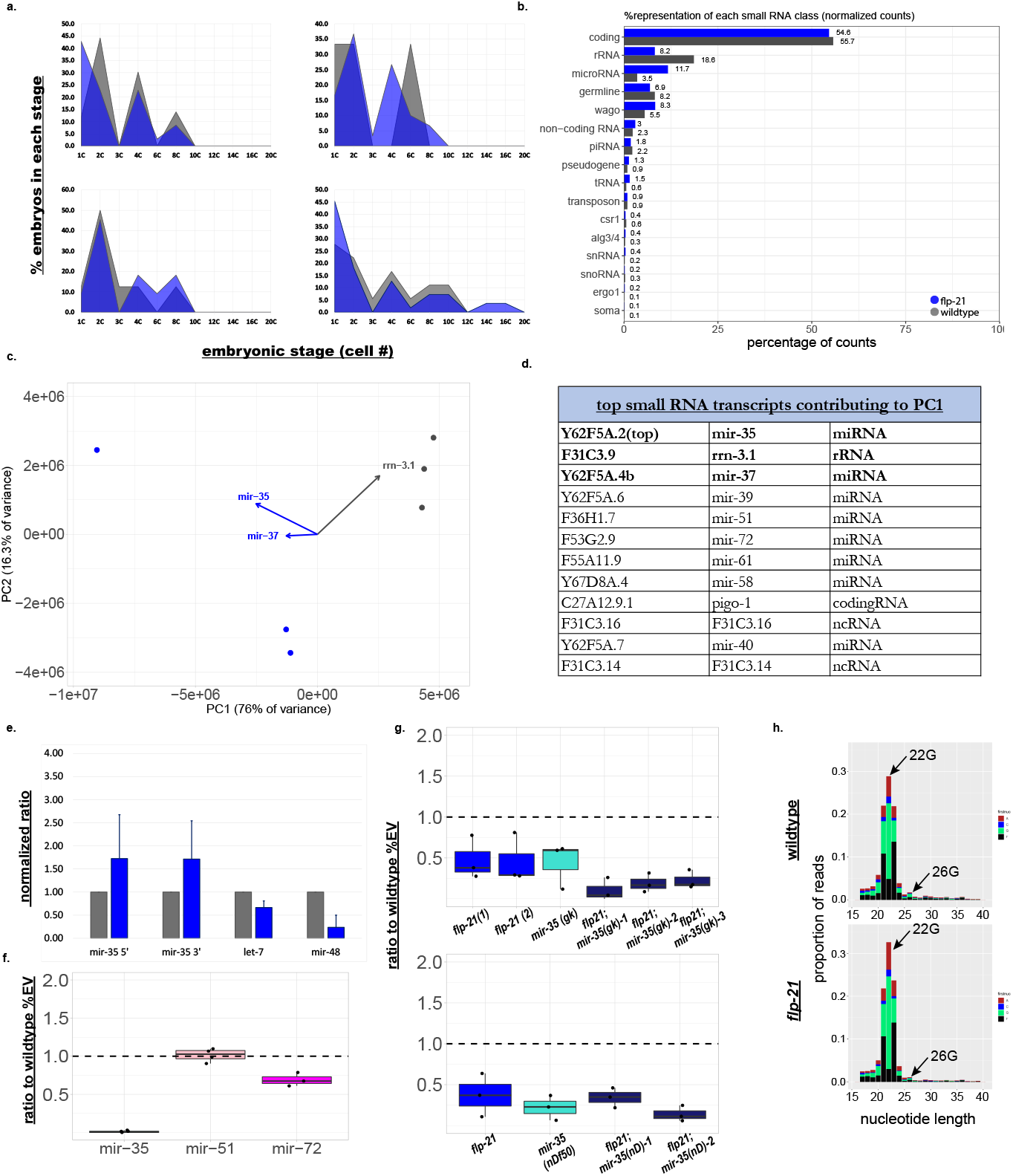
Small RNAs remain largely unchanged in *flp-21* mutants. (a) Embryonic stages obtained for sequenced samples. x-axis represents Embryonic stage (cell number), y-axis represents percentage of embryos for each genotype that fall into specified embryonic stage. (b)Percent representation of each small RNA class in sequencing libraries from *flp-21* and wildtype animals. RPM values used to look at distribution of 16 different small RNA classes. Small RNA classes indicated in RNAi remain unchanged ie piRNAs, CSR1, ALG-3/4, WAGO, and ERGO class small RNAs. (c) First nucleotide distribution of small RNAs graphed as proportion of all small RNA reads between 16-40nt. 22G and 26G siRNA distributions remain largely unchanged (arrow). (d) PCA analysis of wildtype (grey circles) and *flp-21* (blue circles) small RNA samples. Top genes driving clustering of samples labeled and arrows indicate direction of driving change. PC1 on x-axis, PC2 on y-axis. (e) Taqman assay for top small RNA sequencing hits. Data are normalized to internal control and wildtype.

**Figure Supplement 3:**
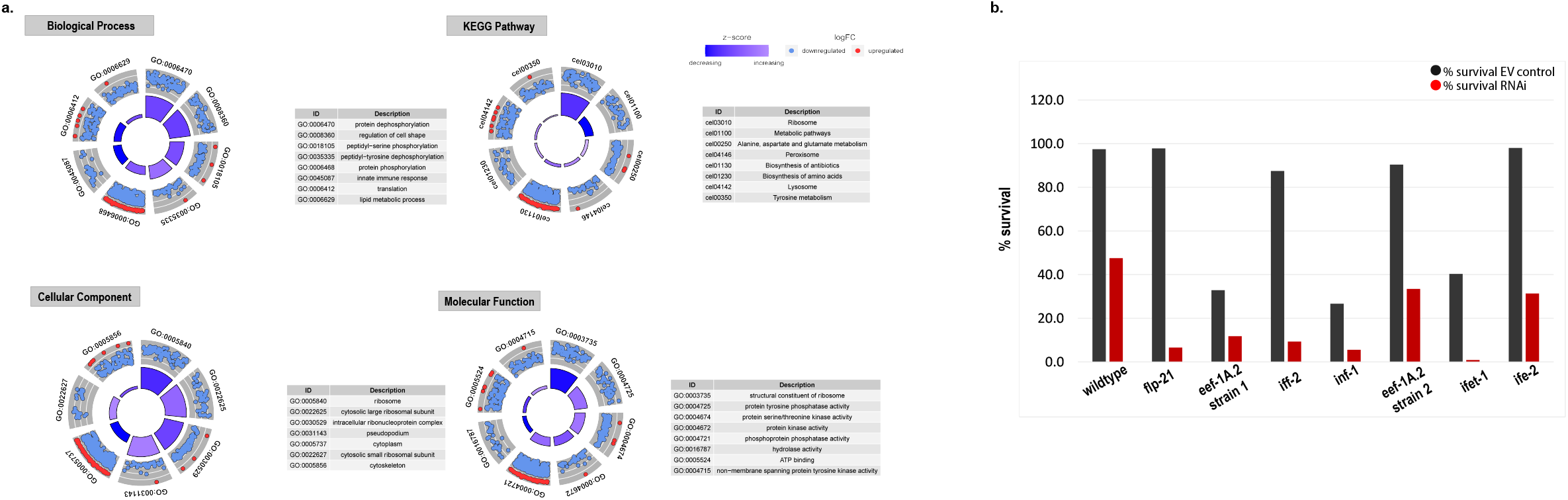
Translational components deregulated in *flp-21* mutants and exhibit *pha-4* RNAi phenotypes. (a) Circle plots of top eight GO terms for each category of analysis (biological function, molecular function, cellular component, and KEGG pathways). Outer circle is scatterplot of log2(foldchange) values for each gene assigned to GO term category. Red dots are upregulated genes, blue dots are downregulated genes. Inner circle bar graphs represent the z-score for each GO term. (b) Raw percent data for EV control survival and *pha-4* RNAi survival data used to Figure 3h. Note low EV survival rates for strains *eef-1A.1* (2), *inf-1*, and *ifet-1*.

## METHODS

### Worm maintenance

Worms were maintained according to Brenner^1^ at 20°C unless otherwise indicated. Strains used supplied in TABLE 1.

**TABLE 1.**
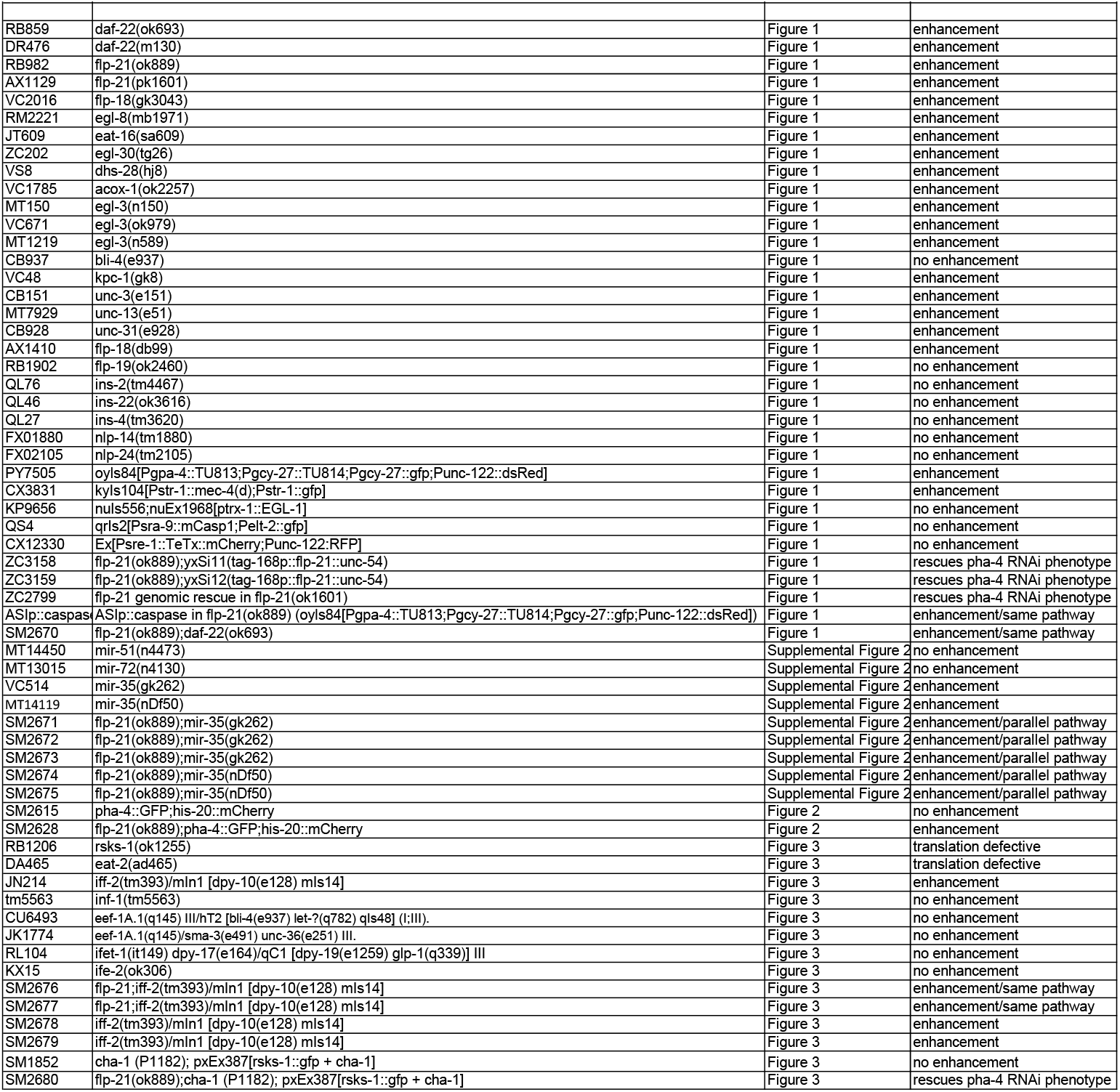

### RNAi

RNAi was conducted according to Wormbook^2^ by feeding L4 larvae - young adult worms bacteria expressing double-stranded RNA for 24 hours (unless otherwise stated). RNAi plates were generated according to Von Stetina, et al.^3^ as follows: Empty Vector(EV; L4440 bacteria) and *pha-4* (PR244) were grown O/N at 37°C with 50μg/ml of antibiotic carbenicillin. OD600 of 1:10 dilution of culture was checked and then concentrated culture was diluted to OD600 1.5 in LB. Cultures were centrifuged for 6 min at 4.4 rpm, room temperature (RT) and the supernatant was removed. Bacteria were resuspended in 10% of original volume in LB. EV(L4440) and *pha-4* (PR244) cultures were mixed together to dilute the *pha-4* dsRNA i.e. 1:4 or 1:5 of *pha-4* dsRNA bacteria to EV dsRNA bacteria. 250ul bacteria were seeded onto NGM plates and allowed to grow for 2 days at RT. Plates were placed at 4°C for 2 days before use. The dsRNA bacteria clones were from the Ahringer library^4^. The identity of the clones was confirmed by sequencing prior to use. For other RNAi clones, bacteria were grown in 5 ml LB with 5 μl carbenicillin (100 mg/ml) for 6-8 h at 37°C and pelleted at 2500 *g* for 10 min. The bacterial pellet was resuspended in 400 μl 0.5 M IPTG, 30 μl 100 mg/ml carb and 70 μl nuclease-free water. NGS plates (5 ml) were seeded with 200 μl of resuspended bacterial solution and kept at room temperature for 2 days, then stored at 4C O/N before use.

### Survival Assay

For each RNAi assay, wild-type control worms were evaluated in parallel to mutant test worms. Fourth larval stage (L4) animals were placed on *pha-4* dsRNA plates at 25°C, unless otherwise indicated, for 24hrs (unless otherwise indicated). Embryos were removed from the RNAi plates and transferred to plates with OP50, allowed to grow for 2 days at either 20° or 25°C, and the surviving adult animals were counted. For the statistics, the number of adults after test RNAi was normalized to the survival of EV counterparts (%EV survival). Statistics were performed according to the experimental design: Wilcoxon Rank Sum (1 control to 1 mutant), ANOVA/Dunnett’s (1 control for 2+ mutants), or ANOVA/Tukey’s (epistasis experiments). For the graphing, wild-type (normalized to %EV) for each experiment was set to 1 and mutant %EV survival was graphed as a ratio.

### Pheromone treatment

Generation of conditioned media: Wild-type N2 worms were harvested by washing off 2x 6cm plates with S media and adding to 250ml cultures. These were grown in S media 250 ml cultures (following worm book protocol: substituting sodium citrate for potassium citrate)^5^ fed with a starting pellet of OP50 *E. coli* roughly equal to that from 500mL of culture. Three cultures were started, kept at 25° C and rotated at 180 rpm. One culture was harvested at 3 days, and the second culture at day 5. The remaining culture was fed again on day 5 and then on day 10, and subsequently harvested on day 12. Each feeding was roughly equal in size and done when the cultures began to become less hazy. To harvest each culture the liquid from the culture was pelleted using a centrifuge at 6500 rpm for 20 min. The clarified media was decanted and freeze dried along with the pellet. The pellets were extracted with a 3:1 mixture of MeOH and CHCl3 and the freeze-dried media was extracted with MeOH and EtOH twice for ~100 mL total. Due to the large amount of dissolved salts the extracts were triturated with MeOH twice and filtered through a plug of celite. This was only done with the media extracts and not the extracts from the pellets.

The ascaroside dauer pheromones, asc-C6-MK (C6; ascr#2), asc-DC9 (C9; ascr#3), and asc-wC3 (C3; ascr#5), were synthesized according to previously published methods^6^.

### Pharyngeal pumping rate assays

Pumping assays per performed as previously described^7^. Briefly, young adults that were well fed were examined for their pumping rate per 20 seconds while on either *E. coli* OP50 lawns or pha-4 RNAi bacterial lawns. Numbers of pumps per 20 seconds were counted. For each experiment 5 young adult worms were analyzed. 2-3 days of assays were performed per mutant genotype. Wild type worms were tested in parallel on each experimental day. For *E. coli* OP50 food lawns, worms were placed on food and left for 1 hour before measuring pumping rates. For pha-4 RNAi plates, adult worms were tested during their 24-hour exposure on 1: 4 dilution pha-4 RNAi bacteria. Mean +/-SE, Student T Test. *p<0.05, ** p<0.01, *** p0.001

### Time course for RNAi sensitivity

Forty L4 worms were placed on EV and *pha-4* RNAi plates at 25°C. After each indicated time point, 100 embryos were removed and placed on OP50 plates and adult worms were moved to new RNAi plates. Survival was assayed as indicated above.

### Antibody staining

Antibody staining was conducted essentially according to^3^. Rapidly growing worms and embryos were washed off of plates with dH_2_O at room temperature then pelleted for 1 min at 4000 rpm. Worm pellets were washed 2x with 500ul dH_2_O at RT. Pellets were resuspended in 300μl dH_2_O bleach solution (4.8mL bleach, 1mL 5M NaOH, 4.2mL dH_2_O) and incubated for 3 min with constant vortexing at RT. The bleaching reaction was stopped by adding dH_2_O up to 1.5 ml. Embryos were washed 3x with 1ml of 1X M9 buffer^5^ at RT and spun down for 1 min at 4000rpm. Embryos were pipetted onto slides and allowed to settle. Excess liquid was removed and replaced with 35ul of fixation solution (2%PFA in PBS). Embryos were squished with a coverslip until ~10% burst then placed into a hybridization chamber for 15 min at RT. Slides were placed on dry ice for at least 1 hour. A freeze/crack was performed by popping off the cover slip, and then the slides were plunged into ice cold MeOH for 3 min. Samples were washed 3x in Tris buffered saline + Tween 20 (TBST; 50 mM Tris-HCl, 150 mM NaCl, 0.05% Tween 20, pH 7.6) for 15 min. Blocking non-specific binding was performed for 30 minutes at 15°C with 100ul TNB blocking solution (Trisbuffered saline (TBS; 50mM Tris-CL, pH 7.5), 150mM NaCl) plus 1% normal goat serum). Primary Antibodies were diluted in TNB at indicated concentrations and samples were incubated overnight at 15°C. Slides were washed 3x in TBS for 5 min each. Secondary antibodies were diluted in TNB at 1:200 concentration and 100ul was added to slides. Slides were incubated at 15°C for 1-2 hours. Slides were washed 3x in TBS for 5 min each. 8ul of 1:3 Vectashield (Thermo Fisher Scientific) DAPI:PBS was added to samples and allowed to incubate for 5 min. A coverslip was placed on slides and sealed with nail polish. Antibodies against the following proteins were used: Endogenous PHA-4 (1:1000)^8^; DLG-1 (1:500; Thermo Fisher Scientific MA1-045).

### Worm culture for embryo isolation

Early embryos were collected from adult worms by bleaching for 5 min in 5x bleach solution (see above). Isolated embryos were shaken at 170 rpm at 20°C in Complete S Medium [100 mM NaCl, 5.6 mM K_2_HPO_4_, 4.4 mM KH_2_PO_4_, cholesterol (10 μg/ml), 10 mM potassium citrate, 2 mM CaCl_2_, 2 mM MgSO_4_, and 1× trace metals] without food overnight. Once the embryos hatched and became L1 larvae, concentrated NA22 bacteria were added to the culture. Synchronized embryos were harvested by bleaching after ~64 hours growth when most worms carried one to two embryos, frozen in liquid nitrogen, and stored at −80°C^9,10^.

### Total RNA isolation

Embryo pellets were resuspended 100uL minimal salts solution (M9)^5^ and 1ml of TRIzol^®^ Reagent (ThermoFisher 15596026) then snap-frozen in liquid N2 and stored at −80°C until all samples were collected. Tubes were placed in a 37°C heat block to warm just until thawed, with intermittent vortexing. Once samples were barely thawed, they were snap frozen in liquid N2 creating one freeze-thaw cycle. This was repeated 4 times. Debris was cleared by centrifuging samples for at 12000g for 10 min at 4°C. Supernatant was transferred to a new tube and incubated for 5 min at RT. 100ul of BCP (1-bromo-3-chloropropane) was added and samples were vortexed for 15 seconds. Samples were incubated for 2-3 min at room temperature and spun for 30 min at 12000g at 4°C. The aqueous layer was transferred to a new tube and 500ul of 4° isopropanol with 1ul Glycoblue (Thermo Fisher Scientific) was added to precipitate the RNA. Samples were placed at −20°C overnight. The next day, samples were spun for 25 min at 12000g at 4°C. The RNA pellet was washed with 500ul cold 75% ethanol then spun at 5 min at 12000g at 4°C. The RNA pellet was air dried in a fume hood and resuspended in 20ul of nuclease-free water. Concentration and quality were measured on BioAnalyzer.

### Small RNA isolation

Small RNAs were isolated according to previous work^11,12^. We used 800 ng of total RNA as starting material. 5′ Modifications of RNA were removed with RppH treatment (NEB Thermopol): 40 ul of Nuclease-free water, 5ul 10X NEB Thermopol buffer, 5ul of RNA for 800ng, and 8ul RppH (5,000 units/ml). Samples were mixed, and then incubated for one hour at 37°C. 1 μl of 500mM EDTA was added and reaction stopped by heating at 65°C for 5 minutes. 1:1 mix of phenol:chloroform was added, and samples were transferred to phase-lock tubes and spun at 14000 rpm at 4°C. The aqueous phase was transferred to new phase-lock tube and 100 ul chloroform was added. Samples were spun for 5 min at 14000 rpm at 4°C and the aqueous phase was transferred to new tube. For precipitation, 2.5X volumes 100% ethanol and 1 μl glycogen were added, and samples were incubated overnight at −80°C. The following day, the samples were spun at 14000 rpm for 30 min and washed with 75% EtOH. All residual EtOH was suctioned off and samples were allowed to air dry for 2 minutes, then resuspended in 6 ul RNase free water. To generate libraries, we used NEBNext^®^ Multiplex Small RNA Library Prep Set for Illumina^®^(Set 1) (catalog#E7300S,NEB) (as described^13^). Samples were multiplexed using barcodes that carried Ilumina Truseq adaptors. Size selection for small RNAs of size 16-40nt was done on 5% polyacrylamide gel and extraction of libraries was performed via manufacturers’ recommendations. Quality of samples were analyzed on a BioAnalyzer. Sequences of standards from synthetic construct for Aequorea victoria partial *gfp* gene for GFP (GenBank: LN515608.1).

### Sequencing analysis

Libraries were subjected to single-end sequencing with 50 base-pair read length on an Illumina HiSeq 2000 platform. Data processing and analyses were performed using the statistical programming environment R. Illumina TruSeq universal primer sequence (AGATCGGAAGAGCACACGTCTGAACTCCAGTCAC) was trimmed using CutAdapt. mRNA and small RNA libraries were mapped to the C. elegans genome (WBcel235) using Burrows-Wheeler Alignment Tool (BWA) and processed with SAMtools^14^. MicroRNAs and piRNAs were identified based on their mapped coordinates. The remaining small RNAs were classified as endogenous siRNAs and mapped to either genes, pseudogenes, tRNAs, rRNAs or transposons using *C. elegans* annotation files (wormbase). siRNAs that mapped to genes were further classified into specific endogenous RNAi pathways that are mediated by particular Argonaute proteins including ERGO-1, ALG-3/4, CSR-1, and WAGO RNAi pathways^11,15^–^19^. featureCounts^20^ was used to count all alignments. Multi-read correction was used on all small RNA and mRNA read counts. Small RNA and mRNA read counts from each library were normalized to the total number of mapped reads. A variance stabilizing transformation was utilized before proceeded with downstream analysis. Each library was analyzed for differential expression using Deseq2^21^. A P value < 0.05 and log_2_fold change of 0.67 were used as cutoffs for differential expression. Principal Component Analysis (PCA) loadings were calculated using RPM values. The variance captured in each principal component was extracted and the correlation between each gene and each principal component was determined. The gene contribution projections of the top 6 genes were determined a graphed in Figure 3. GO-term enrichment was analyzed using DAVID and plotted using Goplot^22^.

### qPCR

cDNA was generated from total RNA using Maxima H Minus first strand cDNA synthesis with dsDNase Kit (Thermo Scientific™ K1681) with 1 μg RNA from each sample. RNA was DNase treated for two minutes at 37°C in a preheated thermomixer then chilled on ice after brief centrifugation. The first strand cDNA synthesis reaction was performed in the same tube according to company recommendations, following RT-PCR or RT-qPCR protocols. For qPCR, 1ul of a 1:2 dilution of cDNA reaction was added to qRT reaction. qRT-PCR was carried out with an Eppendorf XX using PowerUp SYBR Green Master Mix (ThermoFisher, A25780). The following program was used for qPCR: 3 min at 95°C (3 s at 95°C, 15 s at 55°C, and 30 s at 72°C) for 40 cycles, followed by a melting curve. Reverse transcription reactions without the enzyme and water served as negative controls. All reactions were done in triplicate and on two biological replicates. All the values were normalized to *act-1* as an internal control as well as to the transcript levels in untreated wild type via ΔΔCt method^23^. Primer sequences used in TABLE 2.

**TABLE 2.**
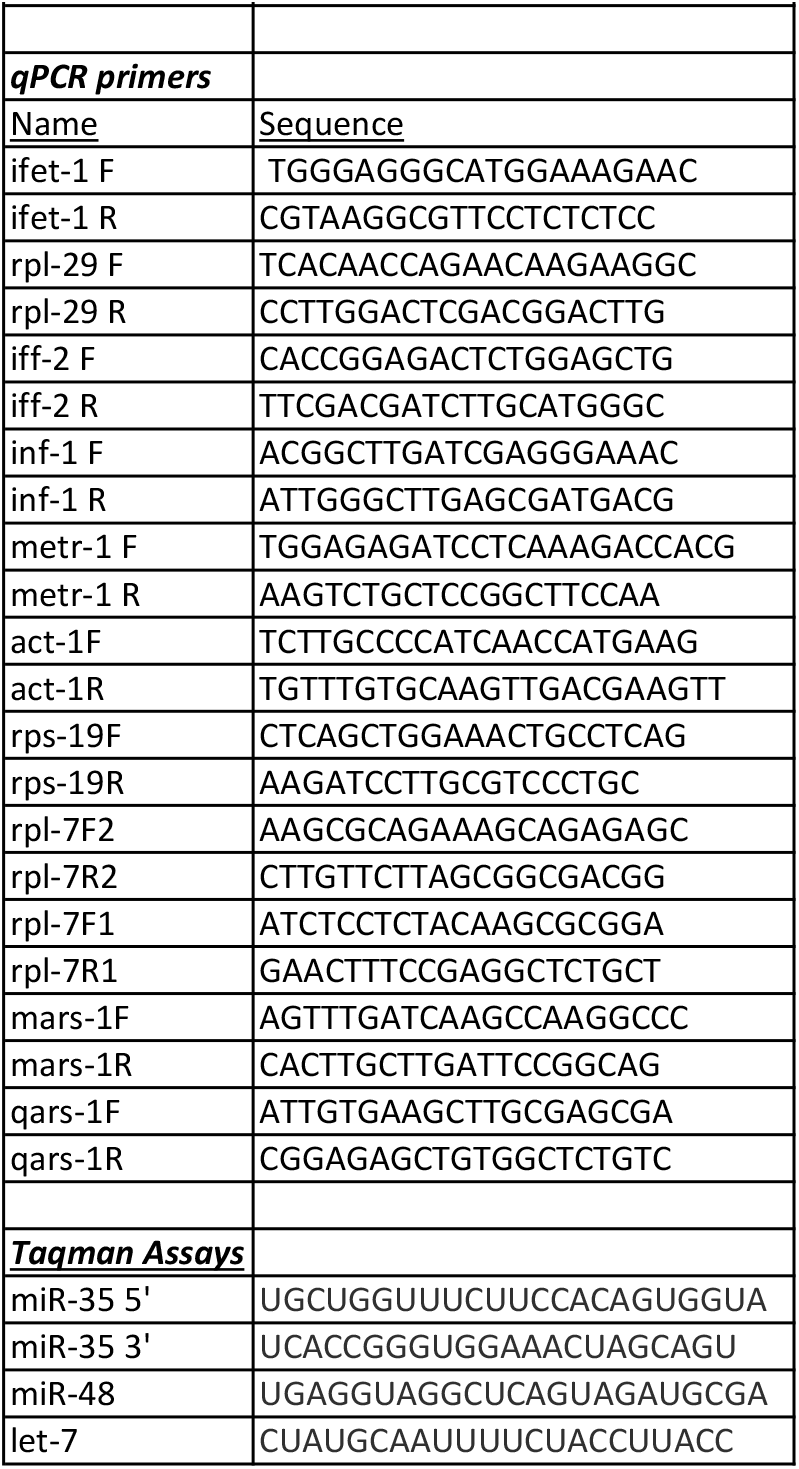

### Taqman Assay

Small RNA quantification was performed as described in previous studies^24^. Levels of miR-35 (5′ and 3′), let-7, and miR-48 were analyzed in wildtype and *flp-21* mutant animals. Assays were performed on 2 independently generated biological samples for both genotypes, and 3 technical replicates were performed for each assay. All smRNA qPCR data were normalized to siR-1 levels. Full small RNA sequences were submitted to Applied Biosystems (TaqMan^®^ MicroRNA Assay 4427975) for design of Taqman assays. Products from each RT reaction were used as template for qPCR using TaqMan^®^ Universal PCR Master Mix II (no UNG; 4440043). Statistical analyses for these qPCRs were performed using T-tests. Probes used in TABLE 2.

### ^35^S metabolic labelling

^35^S metabolic labelling was conducted according to^25^. Worms were synchronized in M9 buffer^5^ and 1000 L1 larvae were plated on fresh OP50 plates and allowed to grow to young adulthood before they produced embryos (~2 days). 10 plates per genotype were grown for each assay plus an additional 10 plates for negative controls. OP50 cultures were grown overnight for a maximum of 12 hrs at 37°C: One culture was grown in LB and one culture with ^35^S methionine at 10μCi /ml (Perkins Elmer). The next day, OP50 cultures were spun down and concentrated 10x. Worms were harvested by washing with S basal^5^ and settling on ice. Worms were washed with 1x S basal, then 1x with S media^5^. Total volume was brought down to 2 ml with 200 μM FUdR (to prevent embryo production; Sigma-Aldrich F0503). 100ul radioactive OP50 was added to test samples and 100ul of non-radioactive OP50 to control samples and then shaken at RT for 5hrs. Samples were then washed with 1x S basal and 1x S media and volume brought down to 2 ml. For negative control samples – 100ul radioactive bacteria was added for less than 1 min with shaking then washed as previously described. 100ul of unlabelled OP50 was added to samples then shaken at RT for 30 min to purge radioactive bacteria out of the intestine. 200ul 1% SDS was added and samples were boiled for 15 min using a heat block set at 100°C and mixed every 5 min for 15 min. Samples were spun at 14K rpm for 15 min and supernatant was removed and placed in new tube. The supernatant was precipitated with 10% trichloroacetic acid (TCA; Sigma-Aldrich T0699) on ice for 1 h. Protein was collected by centrifugation at 14 000 r.p.m. and washed with ice cold ethanol. After air-drying, protein pellets were resuspended in 1% SDS. Protein concentration was measured using Spectrophotometer (NanoDrop) and BCA protein assay kit. 20 ul of sample was taken in duplicate for ^35^S count taken by Beckman scintillation counter. ^35^S incorporation levels were calculated by normalizing ^35^S counts per min, corrected for unspecific background (*t* = 0), to total protein levels. Statistical analysis was done as one-sided, paired Student’s *t*-test on the ^35^S incorporation levels. The relative ^35^S incorporation (used for plotting) was calculated by normalizing the ^35^S incorporation levels of a given mutant to the ^35^S incorporation levels of the control/wild type, which was set to 100.

### smFISH

smFISH was conducted as described in previous studies^26^. Worms and embryos were washed off of rapidly growing plates with dH_2_O at room temperature then pelleted for 1 min at 4000 rpm. Worm pellets were washed 3x with 150μl dH_2_O at RT. Pellets were resuspended in 150μl dH_2_O bleach solution (4.8mL bleach, 1mL 5M NaOH, 4.2mL dH_2_O) and incubated for 4.5 min at 25°C in a thermomixer. The bleaching reaction was stopped by adding dH_2_O up to 1.5 ml. Embryos were washed 3x with 150μl of 1X M9 at RT and spun down for 1 min at 4000rpm. Embryos were pipetted onto slides and allowed to settle. Excess liquid was removed and replaced with 35ul of fixation solution (3.7% FA in 1X PBS/0.05% Triton). Embryos were squished with a coverslip until ~10% burst then placed into a hybridization chamber for 5 min at RT. Slides were placed on dry ice for at least 1 hour. A freeze/crack was performed by popping off the cover slip, and then the slides were plunged into ice cold MeOH for 5 min. Samples were washed 2x in PBS for 5min, and 3x in PBST for 15min. Blocking was performed for 1h at 37°C with 100ul smFISH Hyb buffer –probes(3.7% formaldehyde in 1x PBS/0.05% Triton).

### RNA FISH probes

RNA probes were made by IDT. Primary probe solution (2 ul FLAPY probe + 98 ul Hyb buffer) was added to samples and incubated in a humidity chamber at 37°C for 5 h. Embryos were washed 3x in smFISH wash buffer (10% formamide in 2xSSC/0.5% Triton) for 5 min each. Embryos were washed for 1h at 37°C. Samples were washed 2x more in smFISH wash buffer for 5 min each. 8ul of 1:3 Vectashield Thermo Fisher Scientific) DAPI:PBS was added to samples and allowed to incubate for 5 min. A coverslip was placed on slides and sealed with nail polish. smFISH analysis was performed using the FISHQuant program^27^. smFISH foci were localized in 3D space using Gaussian fitting. Quality control for localized spot is determined by its point-spread function (PSF). Embryos were staged by counting the number of DAPI nuclei.

